# Unique Signatures of Highly Constrained Genes Across Publicly Available Genomic Databases

**DOI:** 10.1101/2024.09.05.611529

**Authors:** Klaus Schmitz-Abe, Qifei Li, Sunny Greene, Michela Borrelli, Shiyu Luo, Madesh C. Ramesh, Pankaj B. Agrawal

## Abstract

Publicly available genomic databases and genetic constraint scores are crucial in understanding human population variation and the identification of variants that are likely to have a deleterious impact causing human disease. We utilized the one of largest publicly available databases, gnomAD, to determine genes that are highly constrained for only LoF, only missense, and both LoF/missense variants, identified their unique signatures, and explored their causal relationship with human conditions. Those genes were evaluated for unique patterns including their chromosomal location, tissue level expression, gene ontology analysis, and gene family categorization using multiple publicly available databases. Those highly constrained genes associated with human disease, we identified unique patterns of inheritance, protein size, and enrichment in distinct molecular pathways. In addition, we identified a cohort of highly constrained genes that are currently not known to cause human disease, that we suggest will be candidates to pursue as novel disease-associated genes. In summary, these insights not only elucidate biological pathways of highly constrained genes that expand our understanding of critical cellular proteins but also advance research in rare diseases.

## Introduction

The advent of large-scale population databases has revolutionized the field of genetics by providing a rich resource for gene and variant-level data^1–3^. Such databases catalog the frequency and distribution of genetic variations across diverse populations, offering a powerful tool for interpreting human biology, and linking genes and their variants with human diseases^4–7^. The genome aggregate database (gnomAD) is one of the most extensive and widely used publicly available collections of population variants from harmonized sequencing data. The gnomAD gene constraint metric for variants is the Z-score^1^, where a high Z-score indicates that a gene is intolerant to variation. Knowledge of a gene’s constraint can inform if genetic variants in the gene are more likely to have a deleterious impact. Recent enhancements to the Z-score metrics in the gnomAD database include loss-of-function (LoF) and missense scores, which can be used to assess complex genotype-phenotype correlations – dependent on mutation type and location^8^.

Evaluating constrained genes as a cohort can provide insights into molecular pathways, the critical role of certain proteins, and facilitate novel gene discoveries^2,6^. Recent literature has emphasized loss-of-function variants for the use of gene discovery due to their high constraint in large population data^1,7^. A thorough evaluation of large-scale public databases such as gnomAD for genes that are uniquely LoF, missense, and both LoF and missense constrained can provide potentially high yield insights into many aspects of cellular functions and human diseases.

We performed a comprehensive evaluation of the gnomAD database to determine highly constrained genes for LoF, missense, and both LoF and missense variants, and compared them with distinctly non-constrained genes to provide critical insights about their chromosomal distribution, size, levels and type of tissue expression, specific cellular functions, and molecular pathways. Those highly constrained genes known to be associated with human diseases provide interesting patterns regarding inheritance, protein function among others, while those yet to be linked should be prioritized for human disease gene discovery.

## Methods

### Z-score selection for highly constrained and non-constrained genes

Genetic constraint is defined as the measure of the amount that a genomic region is under negative selection (intolerant to variation)^9^. The gnomAD v4.1.0 (released November 2023) dataset comprises data aligned to the GRCh38 reference sequence (GENCODE v39), from 730,947 exomes and 76,215 whole genomes. Loss-of-function and missense Z-score data were downloaded from gnomAD website (https://gnomad.broadinstitute.org/downloads). Highly constrained genes and non-constrained genes were selected for analysis. To optimize the sample size of gene lists for each constraint group (LoF-C, loss-of-function constrained; Ms-C, missense constrained; LoF/Ms-C, combined loss-of-function and missense constrained; and N-C non-constrained), specific Z-scores criteria were selected which are described in the results section.

### Analogy between highly constrained and non-constrained genes

Gene coordinates were plotted using karyoploteR (version 1.30.0; https://www.bioconductor.org/packages/release/bioc/html/karyoploteR.html) and RStudio (R version 4.4.1; http://www.rstudio.com/) for gene distribution analysis. This analysis identified whether constrained genes showed specific spatial distribution across the genome at the chromosomal level. Tissue specific transcript levels of the selected genes were assessed using the Genotype-Tissue Expression (GTEx) database (https://www.gtexportal.org/home/). We used GraphPad Prism (version 10.3.0) to generate plots depicting the highest tissue expression levels for each gene, facilitating an understanding of their tissue-specific expression profiles.

Additionally, we utilized the Online Mendelian Inheritance in Man (OMIM) database (query date 03/21/2024) to identify constrained or non-constrained genes associated with human diseases using an annotation developed in RStudio. The inheritance patterns and phenotypes of the disease-associated genes were also extracted from OMIM and recorded. A two-sided Fisher’s exact test was performed to evaluate inheritance patterns with a *P*-value of < 0.05 considered statistically significant.

Protein size data of OMIM recognized genes was extracted from the ensembl canonical transcript (https://useast.ensembl.org/index.html). Data was analyzed with GraphPad Prism (Version 10.3.0; GraphPad Software) and expressed as mean ± standard deviation (SD). One-way ANOVA followed by Tukey’s post hoc test was used for multiple-group comparisons. The numbers of samples per group (*n*) and statistical significance for all comparisons are specified in the figure legends. *P* values < 0.05 was considered statistically significant.

### Novel gene analysis

Genes not previously linked to disease in OMIM, henceforth termed novel genes, were analyzed further using human gene mutation database (HGMD; query date 04/03/2024) and ClinVar database (query date 04/05/2024). Novel genes were identified from HGMD using the following criteria: variants in the gene 1) observed in ≥3 instances; 2) from unrelated families; 3) absent in gnomAD; 4) predicted to be deleterious by HGMD; 5) consistent human phenotypes. The pathogenicity of these genes was evaluated using ClinVar with the following criteria: 1) single gene and 2) observation of pathogenic or likely pathogenic (P/LP) variants. Genes not identified by these filtering criteria were recorded as candidate genes not yet associated with human disease.

### Gene Ontology enrichment and gene family analysis

Gene ontology (GO) enrichment analysis of these high-constraint genes was performed in CYTOSCAPE (version 3.10.1)^9^. Specifically, the ClueGO (version 2.5.9) plugin in Cytoscape was utilized to analyze the GO enrichment^10^. For gene family analysis, researchers manually reviewed high constraint genes and cataloged those appearing ≥ 3x in the same gene family.

### Data analysis (statistics and reproducibility)

Figures were developed with GraphPad Prism (Version 10.3.0; GraphPad Software) and expressed as mean ± standard deviation (SD). *P* values were calculated using Fisher’s exact test (two sided) and results less than 0.05 were considered statistically significant. Microsoft office 365 (Version 16.87) was used to make and design the tables.

## Results

Evaluation of loss-of-function (LoF) and missense Z-score data from gnomAD identified over 18,000 genes (Figure 1a). 2,595 and 2,617genes fell within the top 15% of high-constraint Z-scores, while 346 and 348 genes ranked within the top 2% of Z-scores for LoF or missense variants respectively (Figure 1b). We subdivided the highly constrained genes into three categories: 1) Both LoF- and missense-constrained, or LoF/Ms-C (138 genes; top 2% for both LoF and Ms-C) 2) constrained only for LoF and not missense, or LoF-C (63 genes; top 2% for LoF and bottom 15% for missense); and 3) constrained only for missense and not LoF, or Ms-C (35 genes; top 2% for missense and bottom 15% for LoF). One hundred thirty two genes were classified as non-constrained genes or N-C (not constrained for both LoF and Ms-C; bottom 2%) (Figure 1c **and Table 1**).

**Figure 1.**
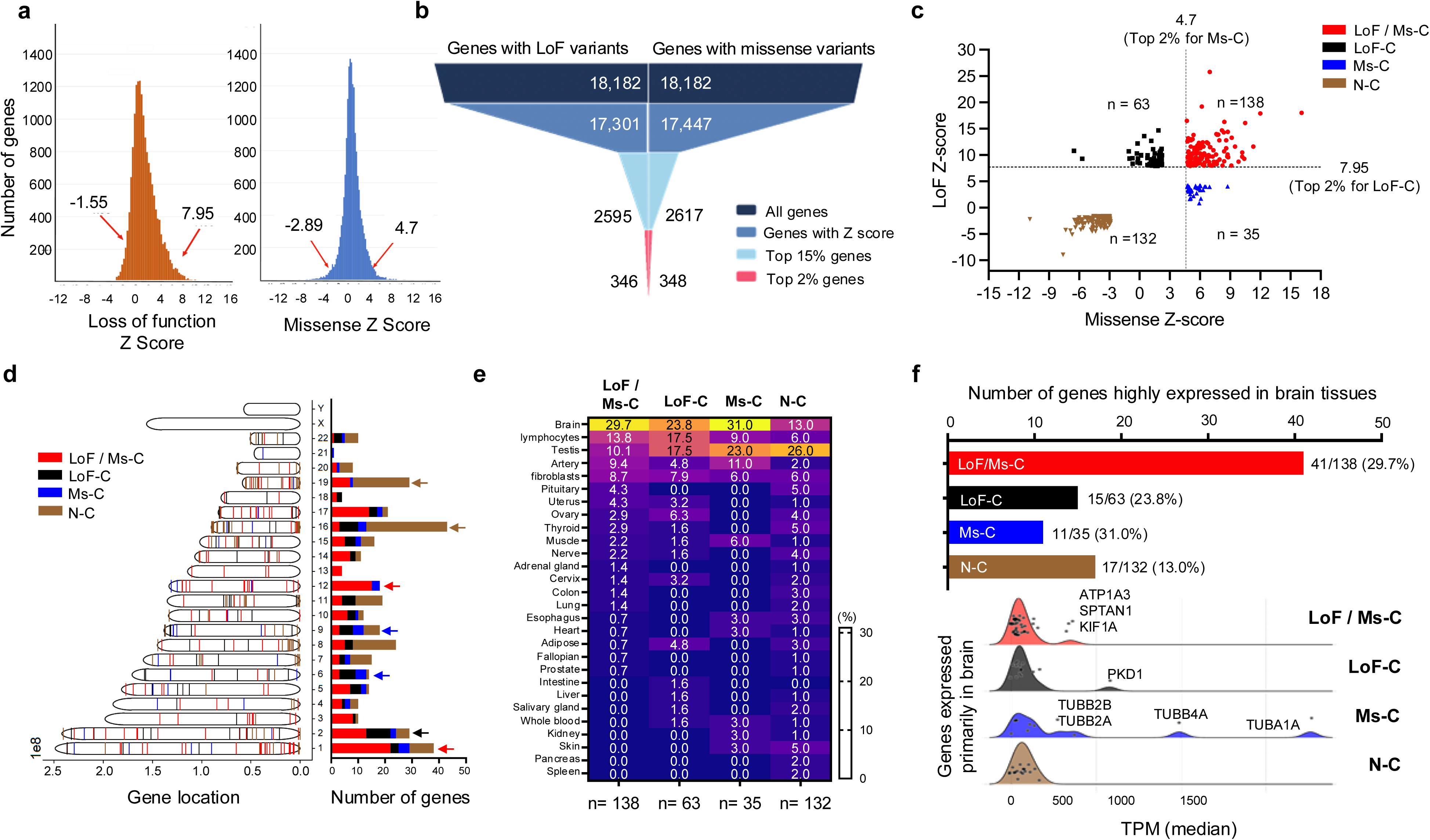
Selection of constraint genes and tissue expression. **a)** Distribution of genes identified with LoF (left) and missense (right) variants according to gnomAD calculated Z-score. Red arrows indicate Z-score cutoff for bottom and top 2% genes respectively (LoF Z-score of < - 1.55 or > 7.95 and missense Z-score < −2.89 or > 4.7). **b)** Visual depiction of selection of top 2% constrained genes from the 18,182 genes identified with LoF (left) or missense (right) variants. Selection required genes with a reported Z-score and the top 15% of constrained genes included 2,595 with LoF variants and 2,617 with missense variants. The top 2% of constrained genes with LoF variants identified 346 genes and the top 2% of constrained genes with missense variants identified 348 genes. **c)** Distribution of constrained and non-constrained genes after grouping of highly constrained genes into three categories (LoF-C, Ms-C, and LoF/Ms-C) based on the missense Z-score to the LoF Z-score provided by gnomAD. **d)** Distribution of genes enriched in the LoF/Ms-C group (red arrows), the LoF-C group (black arrow), the Ms-C group (blue arrows), and the N-C group (brown arrows) on the human chromosomes. **e)** Heatmap of the transcription levels of high-constraint and N-C genes in various tissues through the Genotype-Tissue Expression database. All high constraint gene groups were predominantly expressed in brain tissue (29.5% of LoF/Ms-C, 23.8% of LoF-C, 31.0% of Ms-C). The majority of N-C genes (26.0%) were expressed in the testes and only 13% of N-C genes are expressed in the brain. **f)** Genes expressed in the brain with outliers of TPM > 500 across the highly constrained groups: ATP1A, SPTAN1, KIF1A (LoF/Ms-C group); PKD1 (LoF-C group); and TUBB2A, TUBB2B, TUBB4A, TUBA1A (Ms-C group).

**Table 1:** Distribution of constrained and non-constrained groups based on LoF Z-scores and missense Z-scores and the number of genes in each of the four groups.

|  | **Constrained groups**<br>**(abbreviation)** | **LoF Z scores** | **Missense**<br>**Z scores** | **# of**<br>**genes** |
| --- | --- | --- | --- | --- |
| Constrained<br>(C) | LoF & missense<br>(LoF / Ms-C) | $\geq 7.95$<br>(Top 2%) | $\geq 4.7$<br>(Top 2%) | 138 |
| | LoF (LoF-C) | $\geq 7.95$<br>(Top 2%) | $\leq 2.34$<br>(Bottom 15%) | 63 |
| | Missense (Ms-C) | $\leq 4.21$<br>(Bottom 15%) | $\geq 4.7$<br>(Top 2%) | 35 |
| Non-Constrained | Non (N-C) | $\leq -1.55$<br>(Bottom 2%) | $\leq -2.89$<br>(Bottom 2%) | 132 |

### Chromosomal distribution and tissue expression of high-constraint genes

Analysis of the chromosomal location of high-constraint genes revealed an enrichment of LoF/Ms-C genes on chromosomes 1 and 12, LoF-C genes on chromosome 2, and Ms-C genes on Chromosomes 6 and 9 (Figure 1d**, Table S1**). In contrast, the genes in the N-C group were primarily found on chromosomes 16 and 19. Interestingly, chromosomes X and Y did not contain any genes categorized within the high-constraint and N-C groups.

In addition to chromosomal location, analysis of tissue expression level revealed distinctions between constrained and N-C groups. More than 23% of the genes in constraint groups were highly expressed in the brain, while the highest expression in the N-C group was identified in the testes. Within the N-C group, approximately 26% of genes were expressed in the testes, and only 13% of genes were expressed in the brain (Figure 1e). Exploring the transcription per million (TPM) expression pattern further of the 33 high-constraint testis genes, 31 were also expressed in many other tissues, including the brain, esophagus, muscle, etc. (TPM > 15), and only two were expressed only in the testis. On the other hand, the N-C group shows 12 out of 34 genes (35%) expressed only in testis (*p* = 0.005, Table S2b). Due to the increased expression of the constrained groups in brain tissue, further analysis was performed to evaluate expression of those genes (Figure 1f). Within the constrained groups, a list of genes were identified as outliers with high expression in the brain: *ATP1A, SPTAN1,* and *KIF1A* (LoF/Ms-C group); PKD1 (LoF-C group); and T*UBB2A, TUBB2B, TUBB4A,* and *TUBA1A* (Ms-C group). A detailed evaluation of their transcription levels across tissues is addressed in **Table S2.**

### Comparison of high-constraint genes with known phenotypes in OMIM

Evaluation of the 368 genes (236 constrained genes and 132 N-C genes) revealed 168 identified as known human disease-linked OMIM genes. The number of known disease-causing constrained genes was 144 out of 236 (LoF/Ms-C, LoF-C, and Ms-C groups account for 97, 27, and 20 genes respectively) compared to 24 of 132 genes in the NC group (*p* = 5.5 × 10^−16^, Figure 2a). Among these known OMIM genes, analysis of the inheritance pattern for each constraint group (**Table 2**) revealed unique patterns between groups. A majority of constrained genes were dominant (LoF/Ms-C to N-C *p* < 0.0001, LoF-C to N-C *p* < 0.001, Ms-C to N-C *p* < 0.0001), while the N-C group contains the highest percentage of recessive genes (N-C to LoF/Ms-C *p* < 0.0001, N-C to LoF-C *p* < 0.01, N-C to Ms-C *p* < 0.001) (Figure 2b). A few differences in inheritance patterns were among specific constrained groups, although they were not statistically significant. Specifically, 85% of genes in the Ms-C group were dominant genes, followed by the LoF/Ms-C group containing 80.4%, and the LoF-C group comprising 70.4 % dominant genes, remaining being recessive. In comparison, the N-C group primarily contained recessive genes (58.3%). Furthermore, evaluation of reported phenotypes associated with known disease-causing genes in OMIM (**Table S3)** determined that 87 high-constraint genes affected multisystemic functions and 35 high-constraint genes primarily affected the brain (Figure 2c).

**Figure 2:**
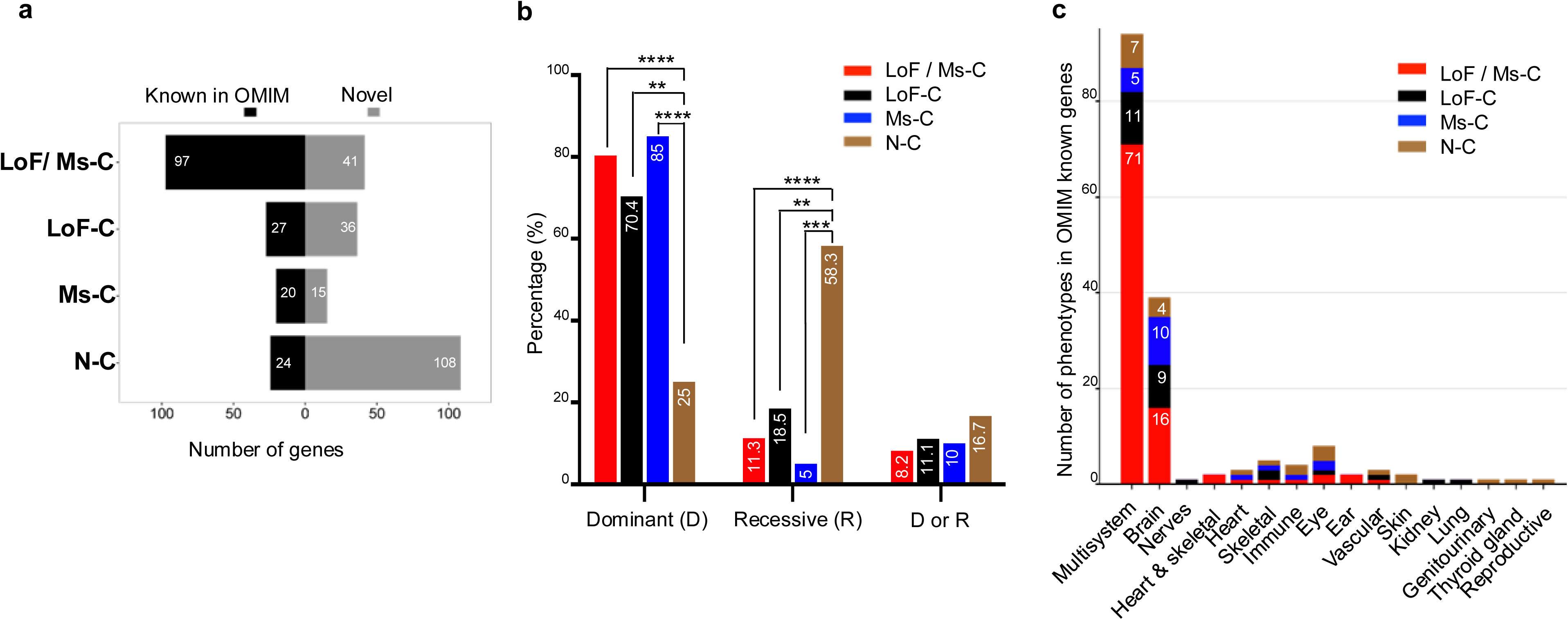
Distribution of constrained and non-constrained genes in relation to association with disease, inheritance and phenotype distribution using OMIM. **a)** Number of disease-associated genes for the constrained group was 144 of 236 (LoF/Ms-C, LoF-C, and Ms-C groups account for 97, 27, and 20 genes respectively) and 24 of 132 for the N-C group. **b)** High-constraint genes were more likely to be dominantly inherited compared to N-C genes, and similarly N-C genes were more likely to be recessively inherited compared to constrained genes. **** *p* < 0.0001, *** *p* < 0.001, ** *p* < 0.01, * *p* <0.05. **c)** High-constraint genes were more likely to be implicated in multisystemic involvement or neurological phenotypes. In comparison non-constraint genes were more likely to affect individual organ systems.

**Table 2:** Inheritance pattern of genes with known OMIM phenotypes in each of the four groups.

| **Groups** | **OMIM-Known** | **Dominant**<br>**(D)** | **Recessive**<br>**(R)** | **D/R** |
| --- | --- | --- | --- | --- |
| **LoF / Ms-C** | 97 | 78 | 11 | 8 |
| **LoF-C** | 27 | 19 | 5 | 3 |
| **Ms-C** | 20 | 17 | 1 | 2 |
| **N-C** | 24 | 6 | 14 | 4 |

Analysis of the protein size (aa) revealed variations across constraint groups. The associated proteins for LoF/Ms-C (2455 ± 3120 aa) and LoF-C (2628 ± 1371 aa) genes were statistically larger than proteins associated with Ms-C (592 ± 323 aa) and N-C (562 ± 512 aa) genes (LoF/Ms-C to N-C *p* < 0.0001, LoF/Ms-C to Ms-C *p* < 0.0001, LoF-C to N-C *p* < 0.001, LoF-C to Ms-C *p* < 0.001) (**Table S1, Figure S1**). There was no statistical significance between the size of LoF/Ms-C and LoF-C gene proteins or between Ms-C and N-C gene proteins.

### Identification of novel genes from high-constraint groups using HGMD and ClinVar

Of the 200 genes not yet linked to human disorder in the OMIM database, 92 were from the constrained groups, and 108 genes were identified in the N-C group (Figure 2a). We reviewed the HGMD and Clinvar databases as well to determine if the 92 novel high constraint genes are implicated in rare or complex disorders, as OMIM entries may not be fully up to date. Five genes in the LoF/Ms-C group were identified in both HGMD and ClinVar, none in other groups. Further evaluation in HGMD alone identified 2, 5, and 1 novel genes classified as disease-causing from LoF/Ms-C, LoF-C, and & Ms-C groups, respectively (**Table 3**). Additionally, evaluation of the pathogenicity of these novel genes in ClinVar only identified 2 additional novel genes in both the LoF/Ms-C and LoF-C, none in other groups. Overall, we found a total of 17 genes that were only present in HGMD and/or ClinVar and not in OMIM (9, LoF/Ms-C; 7, LoF-C; 1, Ms-C; **Table 3**). Following the above analysis, a total of 75 genes in the constraint groups (32, 29, and 14 novel genes within the LoF/Ms-C, LoF-C, and Ms-C groups, respectively) remain to be linked to human disease (**Table S4**).

**Table 3:** Identification of potential novel genes in high-constraint groups using OMIM, HGMD, and ClinVar.

|  | LoF / Ms-C | LoF-C | Ms-C |
| --- | --- | --- | --- |
| gnomAD v4.1 | n = 138 | n = 63 | n = 35 |
| OMIM | 97 (70.3%) | 27 (42.9%) | 20 (57.1%) |
| HGMD & ClinVar | 5 (3.6%) | 0 (0.0%) | 0 (0.0%) |
| only HGMD | 2 (1.4%) | 5 (7.9%) | 1 (2.9%) |
| only ClinVar | 2 (1.4%) | 2 (3.2%) | 0 (0.0%) |
| Unknown | 32 (23.2%) | 29 (46.0%) | 14 (40%) |

### Gene ontology (GO) analysis of high-constraint genes

GO analysis of the 236 genes in high-constraint groups and the 132 genes in the N-C group identified clear clustering of biological processes (Figure 3). Specifically, genes in the LoF/Ms-C group were highly enriched in histone modification (15.6%), ATP-dependent chromatin remodeler activity (9.4%), cardiac muscle cell action potential (8.7%) and post-transcriptional silencing by RNA (5.4%); genes in the LoF-C group were enriched in lysine N-methyltransferase activity (4.8%), histone methyltransferase complex (4.0%), and protein localization to microtubule (4.0%); genes in Ms-C group are associated with a structural constituent of cytoskeleton (18.6%), intracellular bridging (7.1%), and striated muscle development (5.7%). In contrast, genes in the N-C group are enriched in the protein tyrosine kinase binding (4.5%), pathway-restricted SMAD phosphorylation (2.3%), and sensory perception of taste (2.3%).

**Figure 3:**
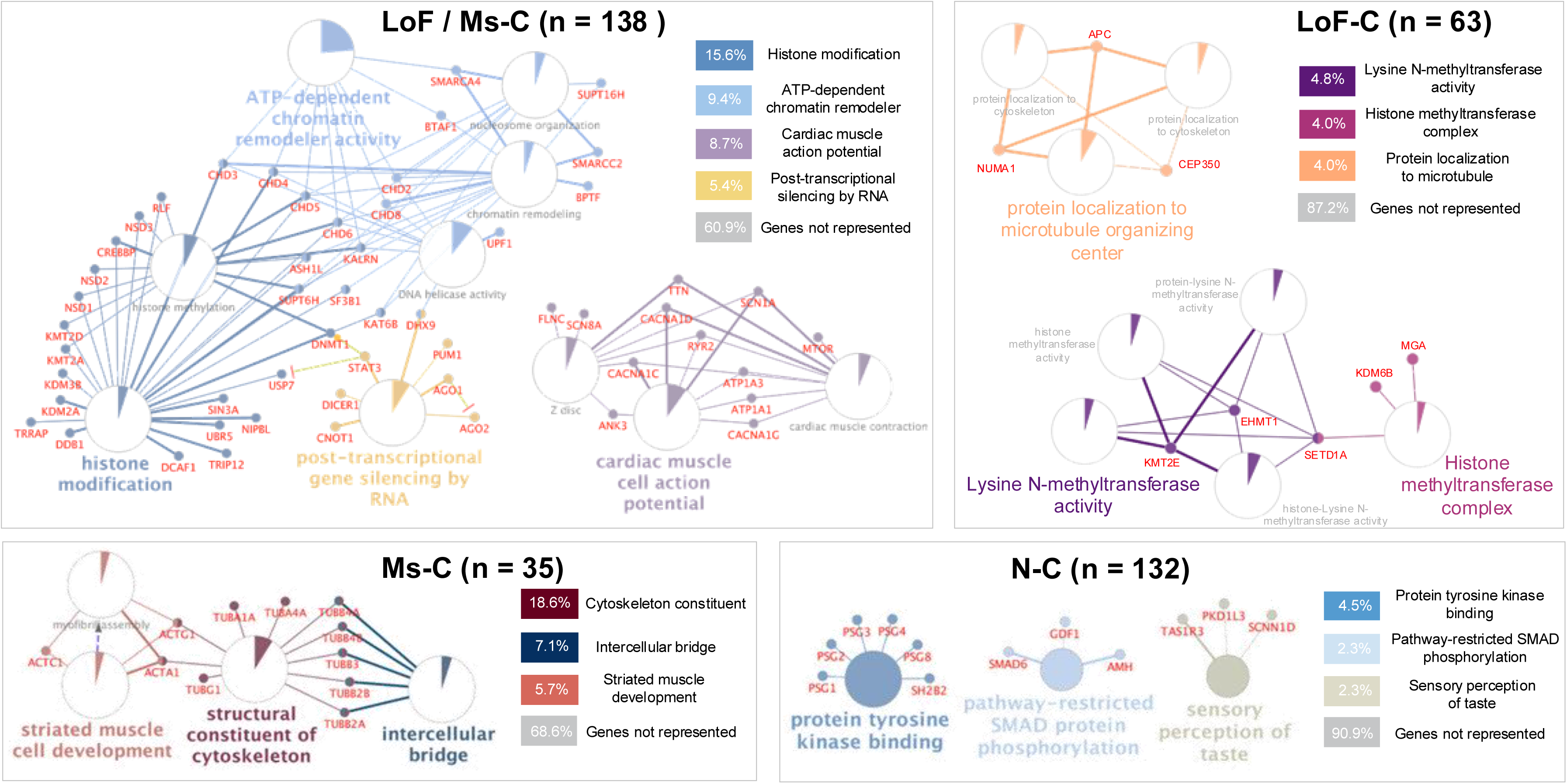
Gene ontology pathway analysis. Gene ontology (GO) analysis of the 236 high-constraint genes and 132 N-C genes identified unique pathway clustering. LoF/Ms-C genes clustered in pathways associated with histone modification (15.6%), ATP-dependent chromatin remodeler activity (9.4%), cardiac muscle cell action potential (8.7%) and post-transcriptional silencing by RNA (5.4%); genes in the LoF-C group clustered in lysine N-methyltransferase activity (4.8%), histone methyltransferase complex (4.0%), and protein localization to microtubule (4.0%) pathways; Ms-C genes were associated with a structural constituent of cytoskeleton (18.6%), intracellular bridging (7.1%), and striated muscle development (5.7%). In contrast, genes in the N-C group clustered in the protein tyrosine kinase binding (4.5%), pathway-restricted SMAD phosphorylation (2.3%), and sensory perception of taste (2.3%) pathways.

In addition to GO analysis, these 368 genes were evaluated according to their gene family classifications (**Table 4**). The LoF/Ms-C group showed a high frequency of genes in families associated with chromodomain helicase DNA, zinc fingers, lysine methylation, Na^+^/K^+^-ATPase, calcium channels, sodium channels, ankyrin, HERC, and ubiquitination genes. In comparison, the LoF-C group exhibited a concentration of zinc finger protein genes, and the Ms-C group was predominantly enriched in structural protein genes such as the tubulin and actin gene families.

**Table 4:** Gene family classifications of the 236 high-constraint genes and 138 non-constrained genes where a gene family was ascertained if ≥ 3 genes in that family were identified.

|  |  |  |  |  |  |  |  |  |  |
| --- | --- | --- | --- | --- | --- | --- | --- | --- | --- |
| LoF / Ms-C | Chromodomain helicase DNA genes | CHD2 | CHD3 | CHD4 | CHD5 | CHD6 | CHD8 |  |  |
|  | Zinc finger genes | ZEB2 | ZFR | ZMYM4 | ZNF236 | ZNF462 | ZSWIM8 |  |  |
|  | Lysine methylation genes | KAT6B | KDM2A | KDM3B | KMT2A | KMT2D |  |  |  |
|  | Na<sup>+</sup>/K<sup>+</sup>-ATPase genes | ATP1A1 | ATP1A3 | ATP2B1 | ATP2B2 |  |  |  |  |
|  | Calcium channel genes | CACNA1C | CACNA1D | CACNA1E | CACNA1G |  |  |  |  |
|  | Ankyrin genes | ANK3 | ANKHD1 | ANKRD17 | ANKRD52 |  |  |  |  |
|  | HERC genes | HECTD1 | HECTD4 | HERC1 | HERC2 |  |  |  |  |
|  | Ubiquitin genes | UBE4B | UBR4 | UBR5 | UBTF |  |  |  |  |
|  | Sodium channel genes | SCN1A | SCN2A | SCN8A |  |  |  |  |  |
|  | Collagen genes | COL1A1 | COL2A1 | COL4A1 |  |  |  |  |  |
|  | Nuclear receptor binding SET domain genes | NSD1 | NSD2 | NSD3 |  |  |  |  |  |
|  | SPT homolog genes | SUPT16H | SUPT5H | SUPT6H |  |  |  |  |  |
|  | UNC-homolog genes | UNC13A | UNC79 | UNC80 |  |  |  |  |  |
| LoF-C | Zinc finger genes | ZFHX3 | ZFHX4 | ZNF292 | ZNF407 | ZNF608 | ZNF638 | ZZEF1 |  |
| Ms-C | Tubulin genes | TUBA1A | TUBA4A | TUBB2A | TUBB2B | TUBB3 | TUBB4A | TUBB4B | TUBG1 |
|  | Actin genes | ACTA1 | ACTC1 | ACTG1 |  |  |  |  |  |
|  | Potassium channel genes | KCNA2 | KCNJ4 | KCNJ6 |  |  |  |  |  |
| N-C | Zinc finger genes | ZCCHC3 | ZFR2 | ZNF467 | ZNF705G | ZNF778 |  |  |  |
|  | Pregnancy specific genes | PSG1 | PSG2 | PSG3 | PSG4 | PSG8 |  |  |  |
|  | Transmembrane genes | TMEM134 | TMEM158 | TMEM88B |  |  |  |  |  |
|  | Solute carrier family genes | SLC16A8 | SLC38A8 | SLC3A1 |  |  |  |  |  |
|  | Olfactory receptors genes | OR4C16 | OR4C46 | OR9G1 |  |  |  |  |  |

The N-C group has a high level of pregnancy-specific and zinc finger genes. Of note, the zinc finger gene family was not enriched in the Ms-C group while it was enriched in the other 3 groups. However, upon evaluation of tissue level expression and GO analysis, we did not identify any unique expression or functional patterns in zinc finger gene families between the three groups.

## Discussion

Evaluation of the genes constrained for LoF/Ms, Ms, and LoF variants in a large population database, gnomAD, provides insights into their unique functional associations and relationship with various Mendelian disorders. This expands upon previous studies’ understanding that constrained genes that are more likely to be associated with disease when mutation occurs^1,2,5,7^. For example, many of these constrained genes are highly expressed in the brain and over 70% of them are implicated in multi-systemic involvement, suggesting their significance in neurobiology and involvement in multiple organ systems.

We identified 236 highly constrained genes and several of these genes have only recently been linked to human disease. For example, *de novo* variants in *MYH10* and *MYCBP2*, two LoF/Ms-C genes, have been recently linked to multi-systemic disorders with significant neurological involvement^10,11^. In addition, our analysis identified *KLHL20* as a Ms-C gene that recently reported *de* novo variants in the motor domain in 14 patients with multisystemic and muscular involvement^12^. The identification of LoF-C gene, *JMJD1C*, was also recently implicated in histone demethylation in a neurodevelopmental and epileptic disorder^13^.

Our analysis identifies 75 highly constrained genes (32 LoF/Ms-C, 29 Ms-C, and 14 LoF-C) that are not yet linked to a human disease using OMIM, HGMD, and ClinVar databases (**Table S4**). We included HGMD and ClinVar for a comprehensive review of disease-causing genes. Novel genes described here are promising candidates as novel human disease genes and for future studies to determine their role in genetic disorders. The discovery of those conditions linked to these genes would provide patients with definitive diagnosis, comprehensive disease management, and potentially lead to the development of new therapies^3,6^.

We identified chromodomain helicase DNA binding proteins and lysine methyltransferases enriched in LoF/Ms-C genes group which suggests involvement in chromatin modification and transcriptional level regulation, crucial for development and cell differentiation^11^. In addition, there were several members of the ubiquitination gene family (*UBE4B, UBR4, UBR5, UBTF genes*) in this category, crucial for cellular degradation pathways and development^6^. GO analysis identified similar pathways between LoF/Ms-C and LoF-C genes in patterns of transcriptional regulation and post-translational modifications, such as histone modification pathways, chromatin modulation through methylation, and gene silencing.

Among the LoF/Ms-C and Ms-C constraint groups, genes associated with solute channels, such as calcium, sodium and potassium channels were identified which suggests a role in cell electrophysiology. In addition, ion channels are known to be critical for numerous transcriptional regulation mechanisms such as promoter interactions, alternative splicing, and post-translational modifications^14,15^. Channelopathies, predominantly caused by genetic variants in ion channel genes, have been implicated in numerous disorders across various organ systems^15^. In the LoF/Ms-C group, several calcium channel genes, such as *CACNA1C, CACNA1D, CACNA1E, CACNA1G*, were identified which have been reported in neurological, cardiac, adrenal disorders^15–17^. Three sodium channel genes, *SCN1A, SCN2A,* and *SCN8A*, were also identified as constrained and are implicated in seizure and migraine disorders and alternating hemiplegia^18–20^. In the Ms-C group, genes encoding potassium channels, *KCNA2, KCNJ4,* and *KCNJ6*, were identified implicated in neuropsychiatric and neurodevelopmental conditions and ADHD^21,22^.

In comparison, the presence of tubulin and actin genes in the Ms-C group suggests their critical role in cytoskeleton structure and dynamics, which are essential for cell shape, division, and transport^10^. Tubulinopathies and actinopathies are critical to nearly all cellular functions and associated with numerous neurodevelopmental abnormalities. Interestingly, tubulinopathies are consistently caused by heterozygous *de novo* missense mutations that are suspected to be gain-of-function and result in abnormal heterodimers^23–26^. Several tubulin genes, such as *TUBA1A, TUBA4A, TUBB2A, TUBB2B, TUBB3, TUBB4A, TUBB4B,* and *TUBG1,* were identified in the Ms-C group (**Table 4**), confirming the understanding that pathologic missense variants are disease causing in highly constrained genes that has been previously undescribed^24,27^. Heterozygous mutations in actin family genes (*ACTA1*, *ACTB*, *ACTG1*) have been implicated in familial cardiomyopathies and a spectrum of myopathies^28,29^.

The gene families identified in the N-C group included taste specific genes and olfactory receptor genes, that are poorly constrained. A gene family whose function is to identify aerosolized chemicals for the function of scent and taste detection would benefit from allowing for more mutations that may be beneficial in a natural selection process if they can identify a chemical that distinguishes a poisonous food or may detect danger sooner.

One of the limitations of this study is the gnomAD database which contains data from a majority of individuals from European ancestry^31^. Including databases with data from more diverse ethnic populations may provide additional insights to constrained genes and associated pathways.

This study presents a methodology to use public, large-scale population databases using a constraint-based approach to identify biologically important proteins, molecular pathways, and prioritizing candidates for rare genetic disorders. The integration of constraint metrics with gene ontology and population databases can provide a better understanding of the pathways disrupted in disease and essential for cellular physiology. As relevant datasets grow, constraint-based approaches will become increasingly helpful and powerful for revealing connections of biological pathways and genetic diseases. Together, these tools and approaches will further gene discovery and expand our understanding of human physiology.

## Supporting information

Supplementary Figure/Legends and Table Titles

## Abbreviations

gnomAD: genome Aggregation Database
LoF/Ms-C: loss of function and missense constrained genes
LoF-C: loss of function constrained genes
Ms-C: missense constrained genes
N-C: non-constrained genes
OMIM: Online Mendelian Inheritance of Man
HGMD: Human Gene Mutation Database
GO: gene ontology
TPM: transcription level per million

## Acknowledgements

We would like to acknowledge Tzofia Drori for her contribution in reviewing and assisting with this manuscript.

## Author contributions

KSA, QL contributed equally to this work and acquired, analyzed, interpreted and visualized data, and drafted the manuscript. SG and MB assisted in drafting and editing the manuscript. SL reviewed and edited the manuscript. MR developed automative methods for data acquisition and analysis. PA conceptualized, edited and supervised the study. All authors reviewed the manuscript.

