## Supplementary Figure/Legends and Table Titles for "Unique Signatures of Highly Constrained Genes Across Publicly Available Genomic Databases"

**Supplementary Figure and Figure/Table Legends**

**Figure S1**

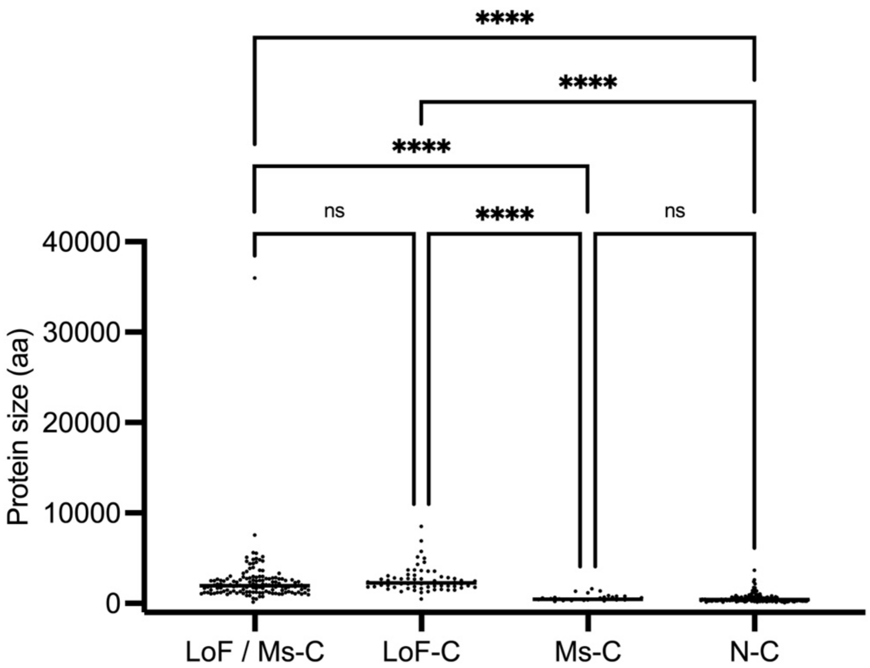


**Figure S1: Reported protein size.** Comparison of constrained and N-C genes revealed LoF/Ms-C (2455 ± 3120 aa) and LoF-C (2628 ± 1371 aa) genes associated with statistically larger proteins than Ms-C (592 ± 323 aa) and N-C (562 ± 512 aa) genes (LoF/Ms-C to N-C $p < 0.0001$, LoF/Ms-C to Ms-C $p < 0.0001$, LoF-C to N-C $p < 0.001$, LoF-C to Ms-C $p < 0.001$). **** $p < 0.0001$, *** $p < 0.001$, ** $p < 0.01$, * $p < 0.05$.

**Table S1:** Reported chromosomal location of 236 constrained and 132 non-constrained genes in GRCh38 and associated protein size (aa) with known UniProt Match.

**Table S2: a)** Tissue specific transcription level of constrained and non-constrained genes.
**Table S2: b)** Transcription levels by constraint groups of genes with TPM < 6, TPM < 10, and TPM < 15.

**Table S3:** Reported OMIM inheritance pattern and known associated phenotypes of disease associated genes.

**Table S4:** List of 75 novel genes currently unlinked with human disease clustered by constraint classification: 32 LoF/Ms-C genes, 26 LoF-C genes, and 14 Ms-C genes.
